# Sterilization of drug-resistant influenza virus through genetic interference: use of unnatural amino acid-engineered virions

**DOI:** 10.1101/2021.12.04.471209

**Authors:** Xuesheng Wu, Zhetao Zheng, Hongmin Chen, Haishuang Lin, Yuelin Yang, Yachao Bai, Qing Xia

## Abstract

The frequent emergence of drug resistance during the treatment of influenza A virus (IAV) infections highlights a need for effective antiviral countermeasures. Here, we present an antiviral method that utilizes unnatural amino acid-engineered drug-resistant (UAA-DR) virus. The engineered virus is generated through genetic code expansion to combat emerging drug-resistant viruses. The UAA-DR virus has unnatural amino acids incorporated into its drug-resistant protein and its polymerase complex for replication control. The engineered virus can undergo genomic segment reassortment with normal virus and produce sterilized progenies due to artificial amber codons in the viral genome. We validate *in vitro* that UAA-DR can provide a broad-spectrum antiviral strategy for several H1N1 strains, different DR-IAV strains, multidrug-resistant (MDR) strains, and even antigenically distant influenza strains (e.g., H3N2). Moreover, a minimum dose of neuraminidase (NA) inhibitors for influenza virus can further enhance the sterilizing effect when combating inhibitor-resistant strains, partly due to the promoted superinfection of unnatural amino acid-modified virus in cellular and animal models. We also exploited the engineered virus to achieve adjustable efficacy after external UAA administration, for mitigating DR virus infection on transgenic mice harboring the 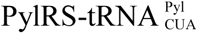 pair, and to have substantial elements of the genetic code expansion technology, which further demonstrated the safety and feasibility of the strategy. We anticipate that the use of the UAA-engineered DR virion, which is a novel antiviral agent, could be extended to combat emerging drug-resistant influenza virus and other segmented RNA viruses.

---

The large-scale use of antiviral drugs for chemoprophylaxis and treatment inevitably imposes strong selection for the evolution of drug-resistant virus. In recent years, high levels of drug resistance have become widespread for human immunodeficiency virus-1 (HIV-1)^1^, hepatitis C virus (HCV)^2^, and influenza A viruses (IAV) that routinely circulate in people^3^. Today, almost all circulating IAV subtypes harboring mutations in M2 protein are resistant to adamantanes^4,5^. In addition, mutations distributed around the active site of neuraminidase (NA), such as E191V, H274Y, R292K, and N294S, endow these viruses with single resistance^6,7^ or multidrug resistance^8,9^ to currently available neuraminidase inhibitors, which increases the risk of a new pandemic and raises an urgent need for developing new anti-influenza countermeasures.

Advances in synthetic biology forward prediction of drug resistant IAV strains emergence^10,11^ and screening for potential antivirals, following an identical antagonism-negates a viral component to interrupt its life cycle^12,13^, yet accelerate the ongoing arm race between virus and traditional drugs which would end up with the spread of evolved drug-resistant virus^14–16^.

Previous studies have demonstrated that defective interfering particles (DIPs) harboring artificial genomic segments can interfere with the replication and packaging of IAV through segment exchange. This represents a paradigm shift of the therapeutic approach by exploiting the biological properties of viral genome. However, the further exploration of this approach is hampered by the low efficiency for the production of DIPs and antiviral treatment^17–19^. The utilization of genetic code expansion^20^ and reverse genetics systems^21^ enables the efficient production of uniform virions containing homogenous defective genomes with artificial amber codons^22^. Whether the uniform virion could be applicable for treatment remains to be shown.

Inspired by genetic interference of DIPs, we proposed that unnatural amino acid (UAA)-engineered virions generated via genetic code expansion would broadly combat virus infection due to a “virus against virus” effect. In this study, we incorporated UAA into viral polymerase complex and drug-resistant protein. We report that the engineered influenza virus could replicate in a controlled way and tested whether it would sterilize normal drug-resistant virus through disseminating defective segments with amber codons. We validated our approach with a massively parallel experimental study to combat a broad-spectrum of influenza viruses and harnessed the optimized UAA drug-resistant virus to reduce symptoms in animal models and protect them from lethal challenges.

## Results

### Genetic manipulation for generation of DR-UAA virus

To test our concept, we first constructed six single-site DR (SDR) IAV strains by introducing a DR-site in PA, NP, or NA protein according to their frequent DR-site mutations^15^ (**Fig. 1a, Extended Data Fig. 1**). The strains included PA-I38T, NP-Y289H, NA-E119V, NA-H274Y, NA-R292K, NA-N294S, and multisite drug-resistant (MDR) strains based on NA mutations, including NA-E119V/H274Y, NA-E119V/R292K, NA-E119V/N294S, NA-H274Y/R292K, NA-H274Y/N294S, NA-R292K/N294S, NA-E119V/H274Y/R292K, NA-E119V/H274Y/N294S, NA-E119V/R292K/N294S, NA-H274Y/R292K/N294S and NA-E119V/H274Y/R292K/N294S, and MDR strains based on different combinations of NA, NP, and PA protein (**Extended Data Fig. 2a**). Next, we systematically explored the relative packaging efficiency (PE) and virus titers of different DR-IAV strains. We found that six SDR strains, six two-site DR strains, and four MDR strains achieved relatively high PE and titers (**Fig. 1b, Extended Fig. 2b**), while others could not be packaged, which might be caused by structural changes of viral proteins after simultaneous multisite mutation, leading to the loss of viral fitness or replication. The successfully packaged DR strains had similar replication kinetics as the wild-type virus (**Fig. 1e, f, Extended Data Fig. 2c, d**). The drug sensitivity test indicated that these DR strains were drug resistant, and MDR phenomena occurred because of introduced multisite DR mutations (**Fig. 1d**).

**Figure 1.**
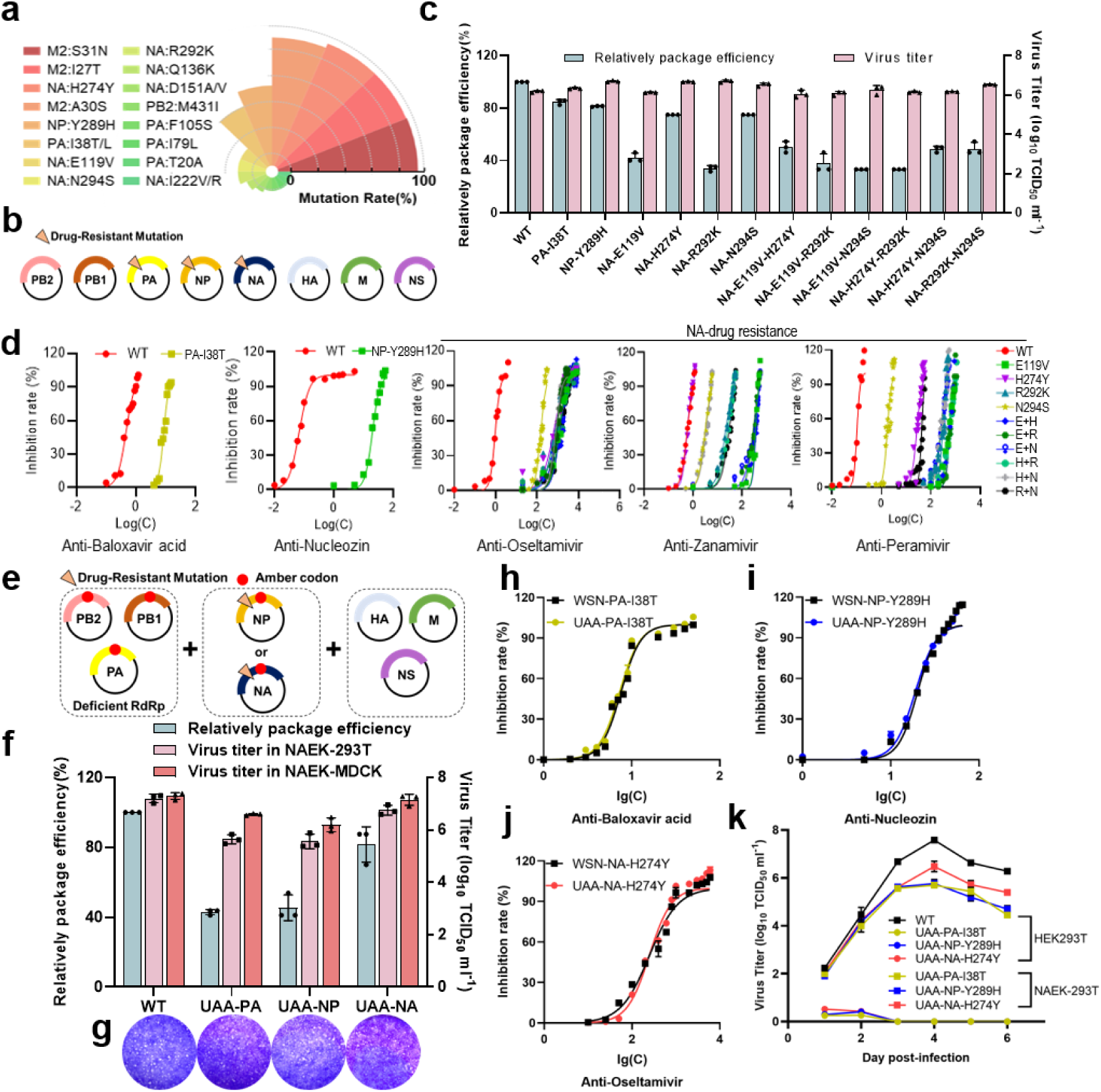
Construction of drug-resistant (DR) UAA virus. (**a**) Statistics of drug resistance mutation sites. (**b**) The introduction of DR sites in PA, NP, and NA segments. (**c**) Relative packaging efficiency and virus titers (median tissue culture infectious dose TCID_50_) of DR strains. Data are presented as mean ± SD (*n*=3). (**d**) Characterization of drug resistance for PA (baloxavir), NP (nucleozin), and NA (oseltamivir, zanamivir and peramivir).

Subsequently, based on our previous experience^22^, we applied reverse genetics to introduce four amber codons (TAG) into the viral genome of successfully packaged DR strains to further generate the DR-UAA virus. These constructs could only replicate in an engineered cell line harboring the PTC-decoding UAA translation machinery, including MbPylRS/tRNA^Pyl^ pairs and unnatural amino acid (NAEK) (**Extended Data Fig. 4**). The amber codon was introduced into the replicase complex (PB2-K33^TAG^, PB1-R52^TAG^, PA-R266^TAG^) and NP-D101^TAG^ or NA-N28^TAG^ to generate DR-UAA virus harboring four amber codons (**Fig. 1e**). The relatively high PE, titers, and plaque formation were observed in UAA-PA-I38T, UAA-NP-Y289H, and UAA-NA-H274Y, among which UAA-NA-H274Y showed the highest PE and virus titer (**Fig. 1f, g**). Similar IC_50_ values of antiviral drug for UAA-PA-I38T, UAA-NP-Y289H, and UAA-NA-H274Y were observed compared with the corresponding DR strains (**Fig. 1h–j**). In addition, the growth kinetic curves and cytopathic effect (CPE) phenotypes of DR-UAA viruses displayed a dependency on UAA concentration in HEK293T engineered cell lines harboring UAA translation machinery (**Fig. 1k, Extended Data Fig. 3a, b**). Collectively, we successfully constructed DR-UAA virus with high virus titers and the capability of drug-resistance which was used for further study.

### Sterilizing effect of DR-UAA virus

The antiviral strategy for the DR-UAA virus was used to produce sterilized viral particles by progeny segment reassortment^23^ when coinfecting with drug-resistant virus (**Fig. 2a**). We co-infected DR-UAA viruses (UAA-PA-I38T, DR-UAA-NP-Y289H, UAA-NA-H274Y, and UAA-IAV without DR as a control) with wild-type WSN and DR strains (WSN-PA-I38T, WSN-NP-Y289H, WSN-NA-H274Y) in MDCK cells, to compare their sterilizing capabilities. The IC_50_ values showed that UAA-NA-H274Y had the best inactivation capability compared to other UAA viruses, which indicated its potential as a therapeutic (**Fig. 2b**). The CPE phenotypes were not observed among all DR-UAA virus groups rather than DR strains (**Fig. 2c**). Collectively, the reassorted progeny harboring the defective gene segment could spontaneously evolve into the sterilized virus, which provides an effective approach for combating infectious DR IAVs.

**Figure 2.**
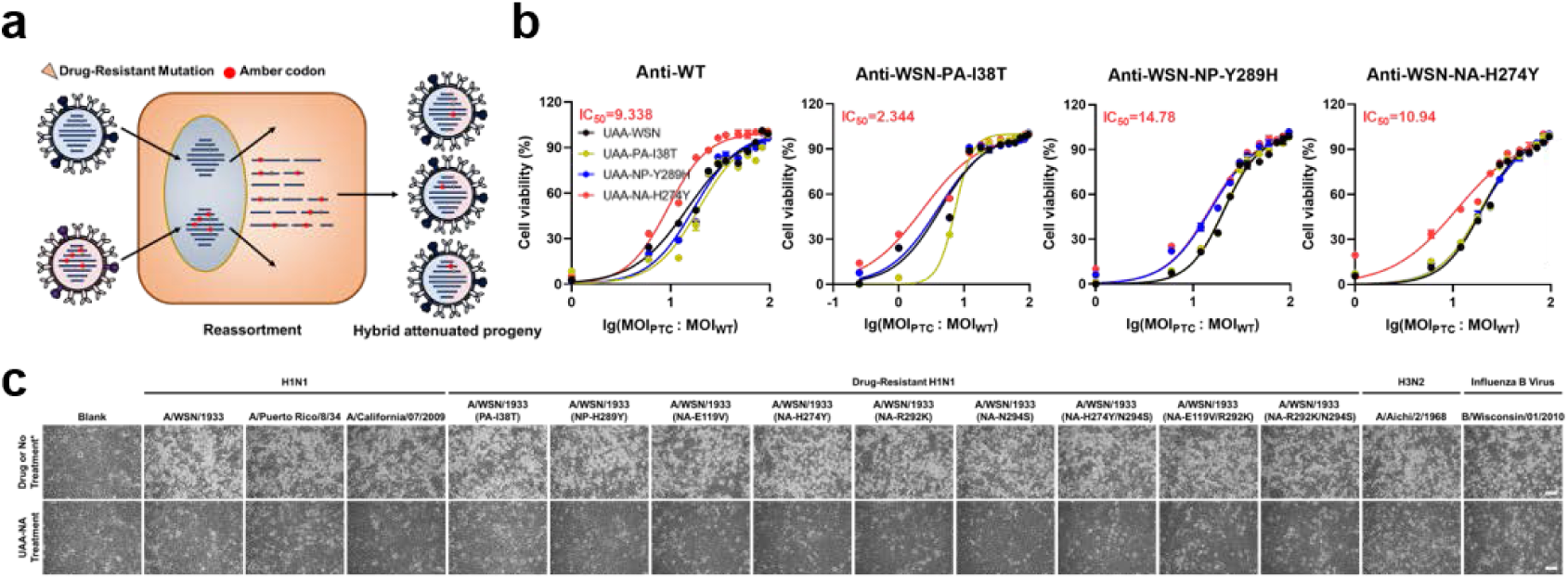
The sterilizing effect of DR-UAA virus. (**a**) Schematic illustration of the sterilized effect of DR-UAA virus combating DR strains after PTC segment reassortment. (**b**) The IC_50_ values of three DR-UAA viruses combating wild-type, DR-PA-I38T, DR-NP-Y289H, and DR-NA-H274Y virus. The common UAA virus was used as a control. (**c**) The cytopathic effect (CPE) phenotype showing the potentially broad-spectrum antiviral ability of UAA-NA-H274Y. Scale bar, 100 μm.

### *In vivo* antiviral efficacy of DR-UAA virus

We performed the antiviral assay in an animal model to investigate whether DR-UAA virus had antiviral ability *in vivo*. Eight BALB/c mice were first intranasally challenged by 1 × 10^5^ TCID_50_ of WSN-NA-H274Y DR strains on day 0, which was followed by treatment with 1 × 10^6^ TCID_50_ of UAA -NA-H274Y strains ± 10 nM oseltamivir on day 1. Phosphate-buffered saline (PBS) and WSN-NA-H274Y **±** oseltamivir were used as controls. Three days after the challenges, three mice from each group were sacrificed for plaque titration of homogenates of the lung and histopathological examination of the lung (**Fig. 3a**). Remarkable improvements of survival rate and body weight were observed in the group of DR-UAA (NA-H274Y) **±** oseltamivir, while all the mice died by day 7 in the WSN-NA-H274Y **±** oseltamivir group (**Fig. 3b, c**). Furthermore, UAA-NA-H274Y **+** oseltamivir showed enhanced potency compared with conditions without oseltamivir, which indicated oseltamivir could further enhance the therapeutic efficacy of DR-PTC (NA-H274Y). Significant amelioration of lung lesions and inflammation were also observed by immunofluorescence and hematoxylin and eosin (H&E) staining in the group of DR-PTC (NA-H274Y) **±** oseltamivir compared to other groups (**Fig. 3d, e**). A significant decrease in RNA copies were observed in the group of UAA-NA-H274Y **±** oseltamivir (**Fig. 3f**).

**Figure 3.**
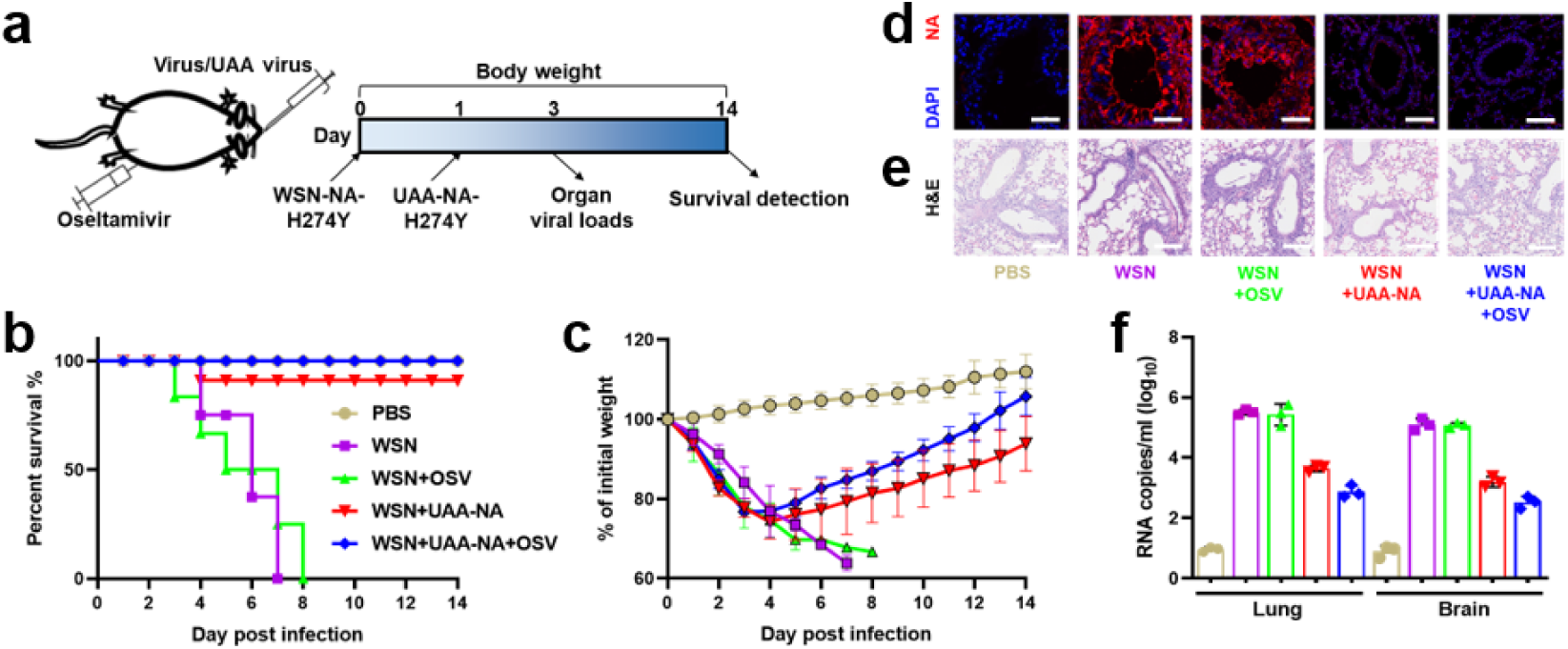
*In vivo* antiviral efficacy of UAA-NA virus. (**a**) Schematic of study design. Mice were first intranasally challenged by DR-NA strains on day 0, which was followed by challenging with UAA-NA strains ± oseltamivir on day 1. Phosphate-buffered saline (PBS), DR strains ± oseltamivir were used as controls. (**b, c**) Body weight changes and survival rates of different groups. (**d**) RNA copies of virus in lung and brain tissue on day 3. (**e, f**) Fluorescent images and H&E staining of day-3 lung tissue in different groups. Scale bars, 200 μm.

### A feasible antiviral strategy based on UAA-controllable virus replication

Next, transgenic mice harboring engineered MmPylRS/tRNA pairs were generated as previously described^24^ and were first intranasally challenged by 1 × 10^5^ TCID_50_ of DR-NA strains and DR-UAA (NA-H274Y) strains **±** UAA. UAA was administrated every day to investigate the functional role of external UAA as a switch in PTC virus replication *in vivo*. PTC virus replication in a UAA-dependent manner was observed by measuring the number of RNA copies and immunostaining analysis (**Fig. 4a, b**). Next, we determined whether the neutralizing effect of DR-UAA virus *in vivo* was UAA-dependent. Transgenic mice were first intranasally challenged by DR-NA strains on day 0, followed by treatment with DR-UAA strains, and different doses of UAA (NA-H274Y) strains + 15 mg UAA on day 1 (low dose: 1 × 10^4^ TCID_50_; middle dose:1 × 10^5^ TCID_50_, high dose: 1 × 10^6^ TCID_50_) (**Fig. 4c**). Remarkable improvements of survival rate and body weight were observed in the DR-UAA virus group, and at different doses of UAA (NA-H274Y) virus **+** UAA, while all the mice died by day 7 in the DR-NA virus group (**Fig. 4c, d**). In the presence of UAA, DR-UAA virus could replicate *in vivo* and even the lowest dose of DR-UAA exerted considerable therapeutic efficacy. These results suggested that the efficacy of DR-UAA virus could be artificially controlled in a flexible manner. Collectively, UAA-NA-H274Y virus displayed significant therapeutic and translational abilities.

**Figure 4.**
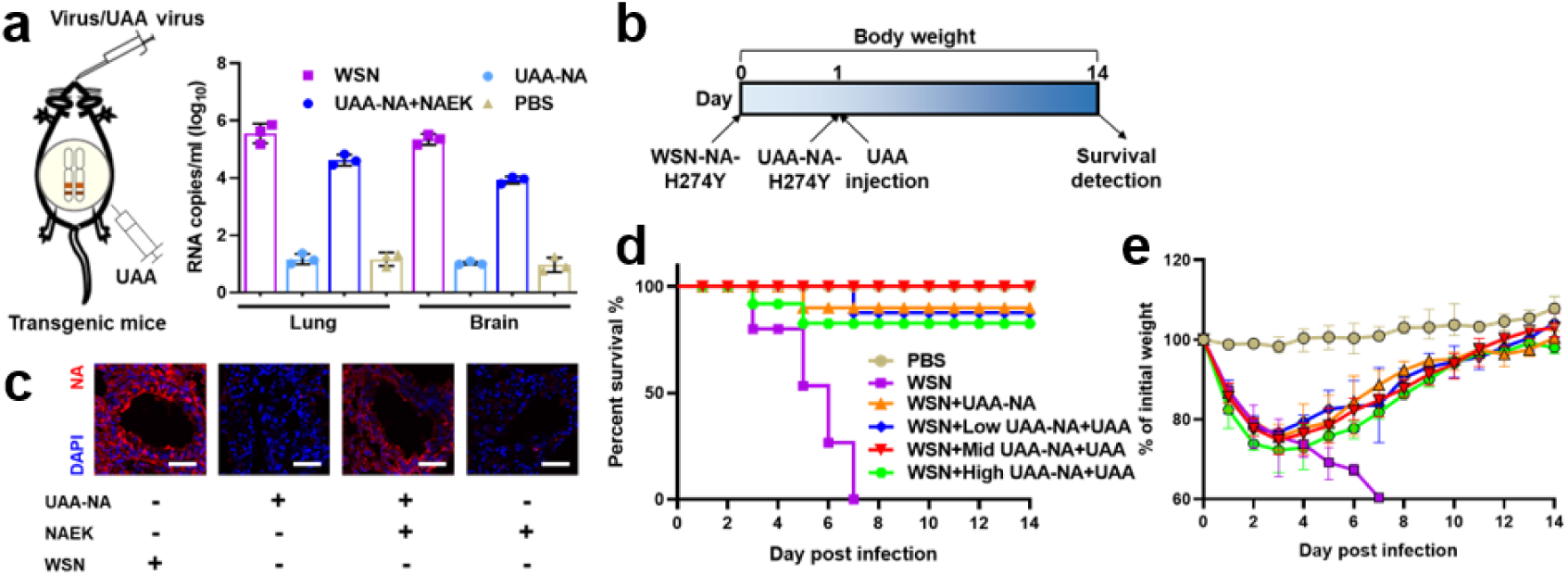
Enhanced efficacy by UAA-controllable virus replication. (**a**) Transgenic mice harboring engineered aaRS/tRNA pairs were generated to achieve regulation of PTC-virus replication with UAA treatment by RNA copies/mL in lung and brain tissue on day 3. (**b**) Immunostaining of NA protein in lung with or without UAA administration on day 3. (**c**) Transgenic mice were first intranasally challenged by DR-NA strains on day 0, which was followed by challenging with UAA-NA strains, or different doses of DR-PTC-NA strains + 15 mg UAA on day 1 (low dose: 1 × 10^4^ TCID_50_; medium dose: 1 × 10^5^ TCID_50_, high dose: 1 × 10^6^ TCID_50_) on day 1. PBS, DR strains ± UAA-NA were used as controls. (**d, e**) Survival rates and body weight changes of different groups.

## Discussion

Influenza virus pandemics are a global health concern and pose a heavy burden on antiviral development. Traditional antivirals, such as antibodies and chemicals, lose efficacy due to antigenic drift or shift and the emergence of drug resistance. This inevitably increases the risk for the next pandemic. We have validated unconventional therapeutic agents, UAA-engineered virions, acting like defective interfering particles in previous studies. The DIPs can interfere with the replication of standard viruses and diminish virus-associated disease with no induction of resistance^25,26^. However, the translational potential of DIPs is limited by low efficiency in its production and the immediate need for high doses in treatment (more than 100 PFU of DIPs are required to neutralize 1 PFU of wild type virus in mice)^17,27^. UAA-engineered virions overcome these obstacles because they are fully infectious like the wild-type virus for a single round infection. They are more competitive than DIPs in their ability to inhibit wild-type replication (the ratio is about 10 to 1 in mice), which results in few safety risks. Moreover, establishment of MDCK-tRNA^Pyl^/PylRS enables the large-scale production of UAA-engineered virions.

This synthetic biology-driven antiviral strategy effectively protects mice from lethal drug-resistant virus challenges, even when given 24 h after infection, which has not been demonstrated by using similar approaches to date. This type of approach also provides novel insights into segment reassortment of influenza viruses. In the *in vitro* neutralization assay, we systematically compare the efficacy of all UAA-engineered virions. We chose UAA-NA for its remarkable efficacy and further investigated UAA-engineered strains carrying different resistant mutations in NA segments. UAA-NA-H274Y exhibited most potential to serve as a broad-spectrum anti-influenza therapeutic, partly due to resembled genome structures^28^ and few segment mismatches^29^ compared with the wild-type virus. No induction of new resistance was observed since it could sterilize the virus by exploiting properties of the viral genome instead of imposing biological stress. Future optimization could focus on generating more segments with amber codons. We have demonstrated the impact of four defective segments at once. More segments will likely be modified simultaneously, especially HA and NS which have high reassorted efficiencies to prevent viral escape. Meanwhile more viral proteins harboring UAA modifications may enable novel properties other than controllable replication^30^.

Another interesting phenomenon was observed when we tested whether UAA-NA-H274Y could work in a simulated clinical scenario. The minimum dose of currently used NA inhibitors, especially oseltamivir, did not inhibit directly, but further enhanced the efficacy of UAA-NA-H274Y to sterilize WSN-NA-H274Y which is resistant to it. Site-specific fluorescent tagging of proteins was used to visualize the behaviors of the viruses. The enhancement to superinfection of UAA-NA-H274Y was attributed to NA inhibitor, which has not been revealed before to our knowledge. However, the mechanism of oseltamivir-mediated superinfection remains unknown, and we propose two potential models. The first is that oseltamivir restores expression of α-2,3- and α- 2,6-linked sialic acids receptors^31^, which are recognized by influenza virus during infection. Nevertheless, oseltamivir could also promote superinfection of WSN-NA-H274Y according to this model, which is contradicted by the result that we found no promotion for the latter. Another model is that the UAA incorporated into the transmembrane domain of neuraminidase influences the translocation of NA^32^ and changes the HA:NA balance in the membrane of UAA-NA-H274Y^33,34^, which leads to the difference in superinfection. Further research is needed to disclose the intriguing mechanism behind this observation. In this study, we exploited a combination of the therapeutic strategy to combat WSN-NA-H274Y in mice and validated the desired protection.

Finally, we designed a proof-of-concept study to answer a question: can we utilize less engineered virions to obtain optimal efficacy by controlling their replication? In transgenic mice with the 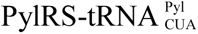 pair, the lethal dose of WSN-NA-H274Y was neutralized by different doses of UAA-NA-H274Y and UAA treatment, in part or in whole, which suggests that restoration of engineered virion replication under UAA control can significantly reduce the dose (the ratio is approximately 1 to 1 in mice). However, before this optimized strategy is put into translational practice, there are two problems that need to be addressed: the formulation of a transient 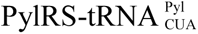 pair delivery system, and identification of the optimal dose window for UAA. When accomplished, this strategy will enable rapid expression of the 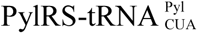 pair and UAA to allow control of a low dose of virion replication in restricted tissues. This should offer fewer safety concerns and lower the cost.

In summary, UAA-engineered virion-based therapeutics represent an alternative approach to the growing problem of drug resistance for influenza virus and might be a more widely applicable approach to use as therapeutics for other drug-resistant viruses.

## Materials and Methods

### Cells and reagents

Human HEK293T and Madin-Darby Canine Kidney (MDCK) cells (ATCC, CRL-2936) were maintained in Dulbecco’s Minimal Essential Medium (DMEM) (Gibco) supplemented with 10% fetal bovine serum (Gibco), 100 units/mL penicillin, and 100 µg/mL streptomycin (Invitrogen). Oseltamivir carboxylate (O126646), Zanamivir hydrate (Z31342), Peramivir (M3222), Nucleozin (A3670), Balxilar (HY-109025) were used in this study.

### Plasmids and constructs

The 12-plasmids allowing the rescue of Influenza A/WSN/33(H1N1) virus by reverse genetics, including the eight influenza viral RNA-coding transcription plasmids (pHH21-PB2, -PB1, -PA, -NP, -HA, -NA, -M, -NS) and the polymerase and nucleoprotein expression plasmids (pCAGGS-PB2, -PB1, -PA and -NP) were kindly provided by Professor George F. Gao and Professor Wenjun Liu from the Center for Molecular Virology, CAS Key Laboratory of Pathogenic Microbiology and Immunology, Institute of Microbiology, Chinese Academy of Sciences, Beijing, China. The wild-type plasmids (pHH21-PA, NP, NA) were used for incorporation of the mutations (PA-R266^TAG^, I38T; NP-D101^TAG^, Y289H; NA-N28^TAG^, E119V, H274Y, R292K, N294S) using appropriate primers and the QuickChangeTM Site-Directed Mutagenesis Kit (SBS Genetech, Beijing, China).

The Tol2 transposon systems transgenes were purchased from Biocytogen (Beijing, China). The methanosarcina barkeri MS Pyrrolysyl tRNA synthetase/tRNA pair (MbPylRS/tRNA) for site-specific incorporation of the unnatural amino acid (UAA) Nε-2-azidoethyloxycarbonyl-L-lysine (NAEK) was developed in-house as previously reported^35^. The PylRS gene driven by CAG promoter was cloned into the Tol2 transposon vector to obtain Tol2-PylRS plasmid. The GFP gene with an amber codon introduced at residue position 39 was expressed under a CMV promoter and cloned into the Tol2-PylRS plasmid to obtain Tol2-PylRS-GFP^39TAG^ plasmid. The pcDNA 3.1 vectors carrying 6 tRNA^Pyl^ copies driven by human H1, human U6 and human 7sk promoters respectively were preserved in our lab. Then it was used as template to arrive the 12 tRNA^Pyl^ copies by Gibson assembly. The 12 tRNA^Pyl^ copies were cloned into the Tol2-PylRS-GFP^39TAG^ plasmid via Gibson assembly to obtain the Tol2-12tRNA^Pyl^-PylRS-GFP^39TAG^ plasmid, which was used for MDCK-tRNA^Pyl^/PylRS stable cell line construction.

### Construction of HEK293T-tRNA/PylRS/GFP^39TAG^ stable cell line

HEK293T cells were used for lentiviral vector packaging and transduction. The cells were cultured in DMEM medium (Macgene, without sodium pyruvate), supplemented with 10% FBS (PAA), and 1 mM nonessential amid acids (Gibco) in 6-well plates until sub-confluent. Then, cells were co-transfected with 0.72 µg of pSD31 transfer plasmid, 0.64 µg of pRSV, 0.32 µg of pMD2G-VSVG and 0.32 µg of pRRE using the transfection reagent Megatran1.0 (Origene). After 6 h, the transfection medium was replaced by DMEM medium containing 3% FBS and 1 mM nonessential amid acids. 48 h post-infection, the lentivirus-containing supernatant was harvested and filtered through a 0.45 µm filter. The resultant dual lentiviruses pSD31-PylRS and pSD31-GFP^39TAG^ were used to integrate PylRS and the GFP^39TAG^ gene into the genome of HEK293T cells. Experiments for stable lentiviral transduction were carried out as follows: HEK293T cells were seeded in a 6-well plate and transduced with lentiviral filtrates in the presence of 8 µg/mL polybrene 24 h later. Then, selection was performed under the pressure of 600 ng/mL puromycin and 200 µg/mL hygromycin until parental cells completely died. The resultant stably transduced HEK293T-pylRS/GFP^39TAG^ cells were transfected with linearized bjmu-12t-zeo plasmid DNA and cultured under the pressure of 200 µg/mL Zeocin until parental cells completely died. In the presence of UAA, the stably transfected cells, HEK293T-tRNA/PylRS/GFP^39TAG^, were then sorted by fluorescence-activated cell sorting (FACS) according to the GFP phenotype and verified by their dependence on UAA for GFP expression.

### Construction of MDCK-tRNA^Pyl^/PylRS for PTC virus

The adherent MDCK cells were maintained in DMEM supplemented with 10% FBS and incubated at 37 °C with 5% CO_2_. For electroporation, MDCK cells were resuspended in Opti-MEM and counted, and 1 × 10^6^ cells were mixed with 15 μg of linearized plasmid mentioned above in medium cup (EC002S). The electroporation was performed using a CUY21EDIT square-wave electropulser (Nepa Gene Co., Ltd) under the following parameters: 250V, poring pulse (5 ms with 50 ms intervals), transfer pulse (50 ms with 50 intervals). After electroporation, the cells were immediately moved from the cup and plated on three 10-cm diameter tissue culture dishes in complete medium with 20% FBS. After 24 h, the plates were washed twice with Opti-MEM and re-fed with complete medium (10% FBS).

To obtain stably transfected MDCK, the cells were first incubated in complete medium with 20% FBS for 72 h and then in complete medium with 4 mM NAEK and 2 μg/mL puromycin. Nearly after 13 days, the MDCK cells containing the transfected gene exhibited a phenotype to form clones. Then the pooled clones were cultured under puromycin pressure. In 15 days, about 80% of the MDCK cells expressed GFP fluorescence, indicating tRNA^Pyl^/PylRS gene was functional to read-through the amber codon introduced in GFP. The cells were sorted by FACS to obtain the monoclonal MDCK-12tRNA^Pyl^/PylRS stable cell line.

### Generation of recombinant influenza A virus

Wild-type influenza A viruses used in this study including Oseltamivir-resistant (NA-H274Y and R292K) influenza A virus A/WSN/33 (H1N1) strains were kindly provided by the Academy of Military Medical Sciences, China. The wild-type and drug-resistant IAV strains were propagated in MDCK cells cultured in DMEM supplemented with 1% bovine serum albumin (Gibco) and 2 µg/mL l-1-tosylamido-2-phenylethyl chloromethyl ketone (TPCK)-treated trypsin at 34 °C for 48 h. The culture supernatant was filtered through a 0.22 µm syringe filter, and viruses were titrated by standard plaque assay procedures.

Mutant viruses were generated by reverse genetic using plasmids as described. For virus rescue, HEK293T cells (or HEK293T-tRNA^Pyl^/PylRS for PTC virus) were co-transfected with 0.2 µg of each of the 12 IAV rescue plasmids per well. Six hours post-transfection, the media was replaced with virus growth media (DMEM, 1% FBS and 2 µg/mL TPCK-treated trypsin). After an additional 24-48 h, we collected media from transfected cells and performed a plaque assay on MDCK (or MDCK-tRNA^Pyl^/PylRS for PTC virus) monolayers to determine viral titer and isolated clones for further growth and characterization.

### Neutralization assays

MDCK cells at 70% confluency in 96-wells plates were incubated with wild-type influenza A virus (WSN) or recombinant DR virus at a MOI of 0.1 for 1 h. Two hours later, the cells were incubated with DR-PTC viruses at different MOI for additional 1 hour. Then cells were put into the incubator and continuously cultured until the cells without any live second DR-PTC virus, followed by cell viability detection using the cell-titer Glu kit (Promega).

### Influenza A virus quantification by 50% tissue culture infective dose

For quantifying all viruses’ stocks, the 50% tissue culture infectious dose (TCID_50_/mL) titers were determined. In brief, 5×10^4^ MDCK or (MDCK-tRNA^Pyl^/PylRS for PTC virus) cells were seeded in 96-well plates the day before infection. The viral samples were serially diluted with DMEM containing 1% FBS (10^3^ to 10^10^) and 2 µg/mL TPCK-treated trypsin and then each of dilution were added in wells separately. The plates were incubated at 37 °C in 5% CO2 for 2-5 days. The cytopathic effect (CPE) was observed under a microscope and virus titer was determined using the Reed-Münch endpoint calculation method^36^.

### Western blot analysis

HEK293T cells were lysed in RIPA lysis buffer (Applygen) supplemented with complete protease inhibitor cocktail (Roche) 48 h after transfection, and cell debris was removed by centrifugation. Tissue was then broken with a homogenizer and removed by centrifugation at 4 °C. Protein was extracted with lysis buffer and quantified by BCA assay (Thermo). A total of 100 µg protein from each sample was boiled with loading buffer, separated on 4%-12% NuPAGE (Invitrogen), and then electroblotted onto a polyvinylidenedifluoride membrane. The membrane was blocked with 5% (v/v) nonfat milk in TBST (50 mM Tris-HCl, 150 mM NaCl, and 0.02% Tween-20, pH 7.5) at room temperature for 1 h and then incubated with rabbit/mouse polyclonal antibodies overnight at 4 °C. Anti-GAPDH protein antibodies (1:1000, sc-365062, Santa Cruz Biotechnology), anti-GFP antibodies (1:1000, 2956T, Cell Signaling Technology), anti-PylRS antibodies (1:1000, sc-40, Santa Cruz Biotechnology, to detect Myc-tagged PylRS) were diluted in TBST containing 5% (v/v) of defatted milk. The membranes were rinsed three times with TBST and then incubated with horseradish peroxidase-conjugated goat anti-rabbit/mouse IgG (1:3000) at room temperature for 1 h. After extensive washing, the protein bands were developed using an enhanced chemiluminescent detection kit (Millipore). The optical bands were visualized in a Fuji Las-3000 dark box (FujiFilm), and band densities were quantified using Quantity One Analysis software. All western blot experiments were done for at least three times in parallel and the representative results were reported.

### Immunofluorescence

The lungs harvested from different mice were fixed in 4% PFA at room temperature for 1 hour, followed by immersing in 30% (w/v) sucrose until submersion before embedding and freezing in the Optimal Cutting Temperature (OCT) compound (Tissue-Tek). Serial 12 µm sections were obtained by cryo-sectioning of the embedded lungs tissue at -20 °C using a cryostat (Leica). Cryosections were blocked with 5% normal donkey serum (Jackson ImmunoResearch) in PBST for 30 min. The sections were incubated with anti-NA antibody (1:500, GTX629696, GeneTex) diluted in blocking buffer at 4 °C overnight. The slides were subsequently incubated with secondary goat anti-rabbit IgG Alexa Fluor 594 (1:400, LifeTech) at room temperature for 1 h. The slides were stained with 0.5 µg/mL Hoechst and mounted in mounting media. Stained sections were photographed under a Nikon Ti-S microscope, and the dystrophin-positive cells were analyzed.

### Histological analysis

The lung tissues of different mice were isolated and fixed in 4% PFA solution for two days, sequentially dehydrated with 70%, 95% and 100% ethanol, and defatted with xylene for 2 h before being embedded in paraffin. The 10-μm-thick section was cut and subjected to hematoxylin and eosin staining (H&E staining) and Masson staining. The slices were observed under optical microscopy, and the histological morphology of different mouse groups was compared.

### Flow cytometry

Until transfected cells were treated with antibiotics to form clones, cells were dissociated into single cell using trypsin/EDTA and analyzed on BD FACSAriaTM (BD Biosciences) with the appropriate filter settings (488 nm coherent sapphire laser for GFP excitation). The front and side scatters were used to identify intact cells and mean background fluorescence from untransfected cells was subtracted from the measured signal. GFP positive cells were harvested for further validation of integrated genes.

### Animal experiments

All animal experiments were performed in accordance with the guidelines of the Institutional Animal Care and Use Committee of the Peking University. Briefly, groups of 4-6-week-old female BALB/c mice (n=8) were anesthetized and intranasally inoculated with 1×10^5^ TCID_50_ of DR-NA-H274Y virus on day 0, then followed by 1×10^7^ TCID_50_ of DR-PTC (NA-H274Y) strains **±** 10 nM Oseltamivir on day 1. PBS, WSN-NA-H274Y **±** Oseltamivir were used as controls. Three days after challenges, three mice from each group were sacrificed for plaque titration of homogenate of lung and histopathological examination of the lung.

All Mice were monitored daily for body weight and survival rate. Two weeks post-inoculation, five mice of each group were sacrificed. The sera were used to test their immunogenicity by ELISA and organs (e.g., brain, intestine and skeletal muscle) were collected to detect viral titer by qRT-PCR.

### Quantitative Real-Time PCR analysis

Total RNA harvested from cells or mice tissues infected with distinct viruses or vaccine was extracted by using Trizol reagent (Invitrogen, CA, USA). RNA (1 μg) was subjected to RT-PCR in accordance with the protocol provided by Promega. The transcripts were quantitated and normalized to the internal GAPDH control. The primers used in the experiments are listed in **Extended Data Table 3**. The PCR conditions were 1 cycle at 95 °C for 5 min, followed by 40 cycles at 95 °C for 15 s, 60 °C for 1 min, and 1 cycle at 95 °C for 15 s, 60 °C for 15 s, 95 °C for 15 s. The results were calculated using the 2^-△△CT^ method according to the GoTaq qPCR Master Mix (Promega) manufacturer’s specifications.

### Statistical analysis

Data are presented as mean ± SD. In most circumstances, three replicates were used for each group unless otherwise noted. The statistical significance between two groups was determined by student’s t test. For three or more groups, one-way or two-way ANOVA with Tukey’s multiple comparisons test was performed. Homogeneity of variances among groups was confirmed using Bartlett’s test. Conformity to normal distribution was confirmed using the Kolmogorov-Smirnov test. All tests were two-sided and performed in GraphPad Prism 8. A probability of p < 0.05 was considered as statistically significant. For annotations, * p<0.05; ** p<0.01; *** p<0.001; **** p<0.0001.

### Reporting summary

Further information on research design is available in the Nature Research Reporting Summary linked to this paper.

## Data availability

The authors declare that all data supporting the results in this study are available within the paper and its **Extended data figure/table legends**.

## Acknowledgments

We thank all the lab members’ contribution to this work. Q. X. was supported by the National Science and Technology Major Projects for “Major New Drugs Innovation and Development” (No. 2018ZX09711003-001-003, 2018-2021).

## Author contributions

Q. X. conceived the idea and designed the experiments. X. W., Z. Z. and H. C. performed the experiments and organized the results, X.W., Z. Z., H. L, H. Z., and Q. X. analyzed the data, prepared the figures and wrote the manuscript. All other authors participated in some of the experiments, results or discussion. Q. X. supervised the study. X. W., Z. Z. and H. C. contributed equally to this work.

## Additional information

**Supplementary information** is available for this paper at the supplementary materials.

**Correspondence and requests for materials** should be addressed to Q. X.

---

**Extended Data Figure 1.**
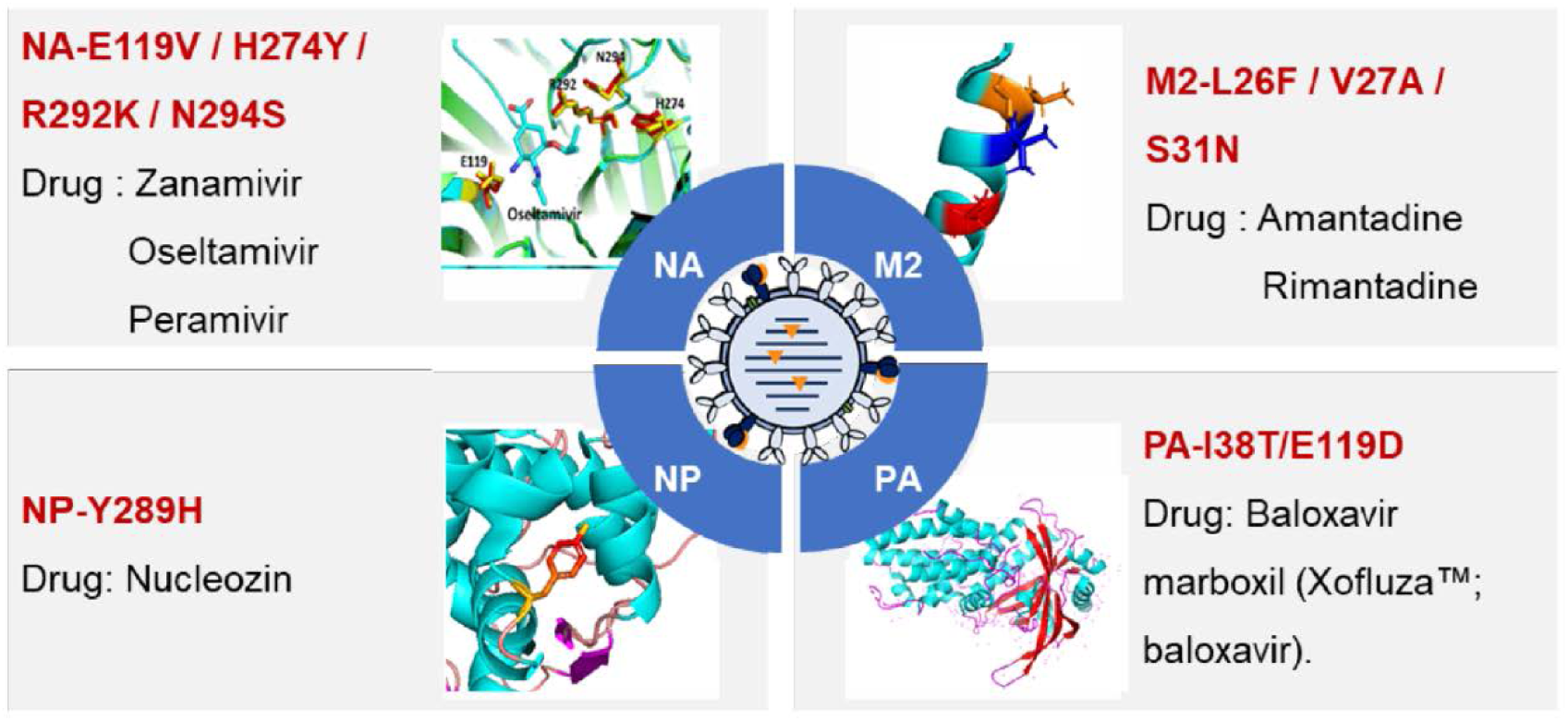
Overview of drug resistance and binding domain in different segments, including NA, NP, and PA. Table shows the combination of different DR sites in different segments (**Related to** **Fig. 1a**).

**Extended Data Figure 2.**
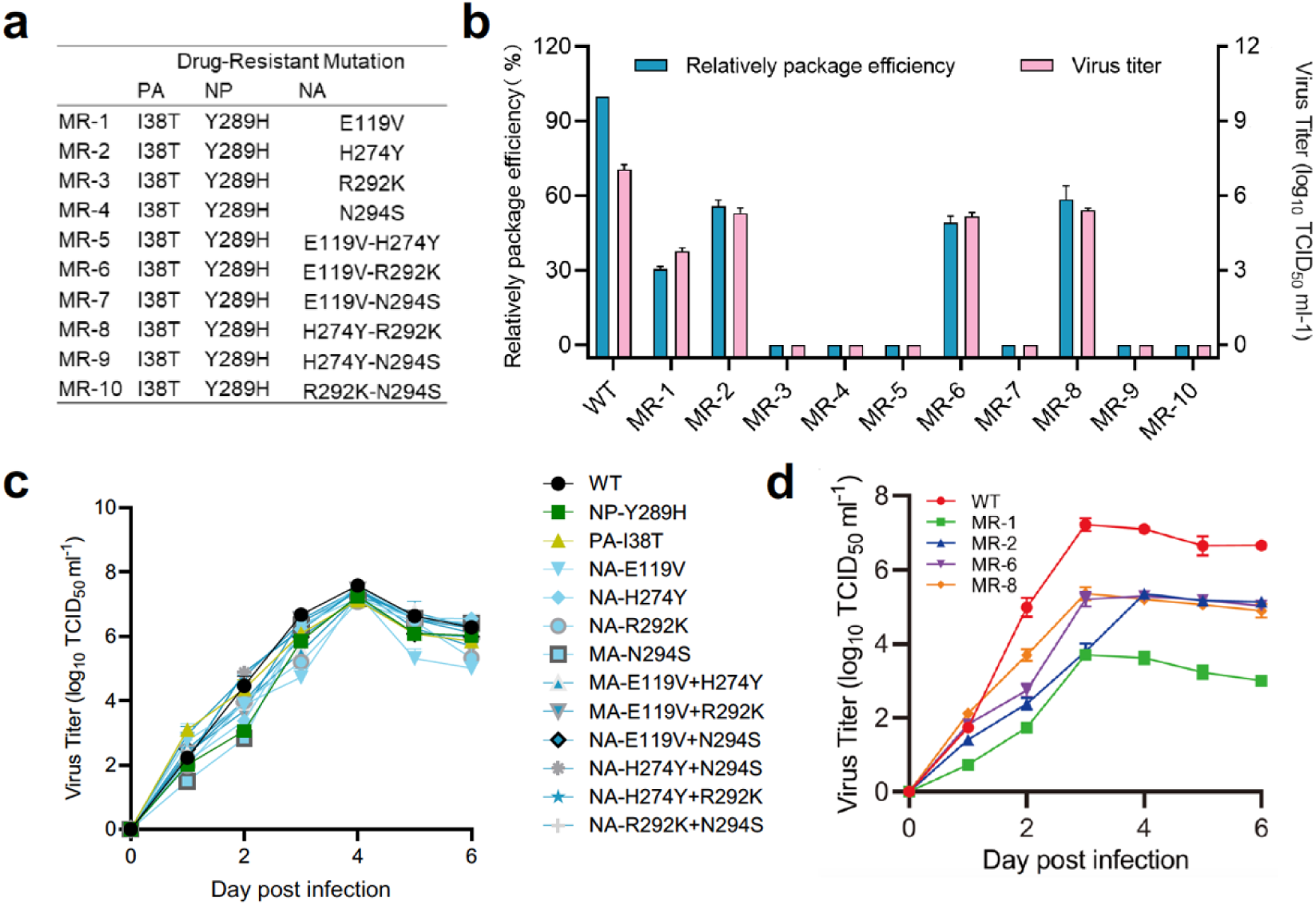
Construction of multidrug-resistant UAA virus. (**a**) The table shows the combination of different DR sites in different segments. (**b**) The relative packaging efficiency and virus titers of all the multidrug-resistant UAA strains. Data are presented as the mean ± SD (*n*=3). (**c, d**) Growth kinetic curves of DR-UAA virus and multidrug-resistant strains (**Related to** **Fig. 1c**). Data are presented as the mean ± SD (*n*=3).

**Extended Data Figure 3.**
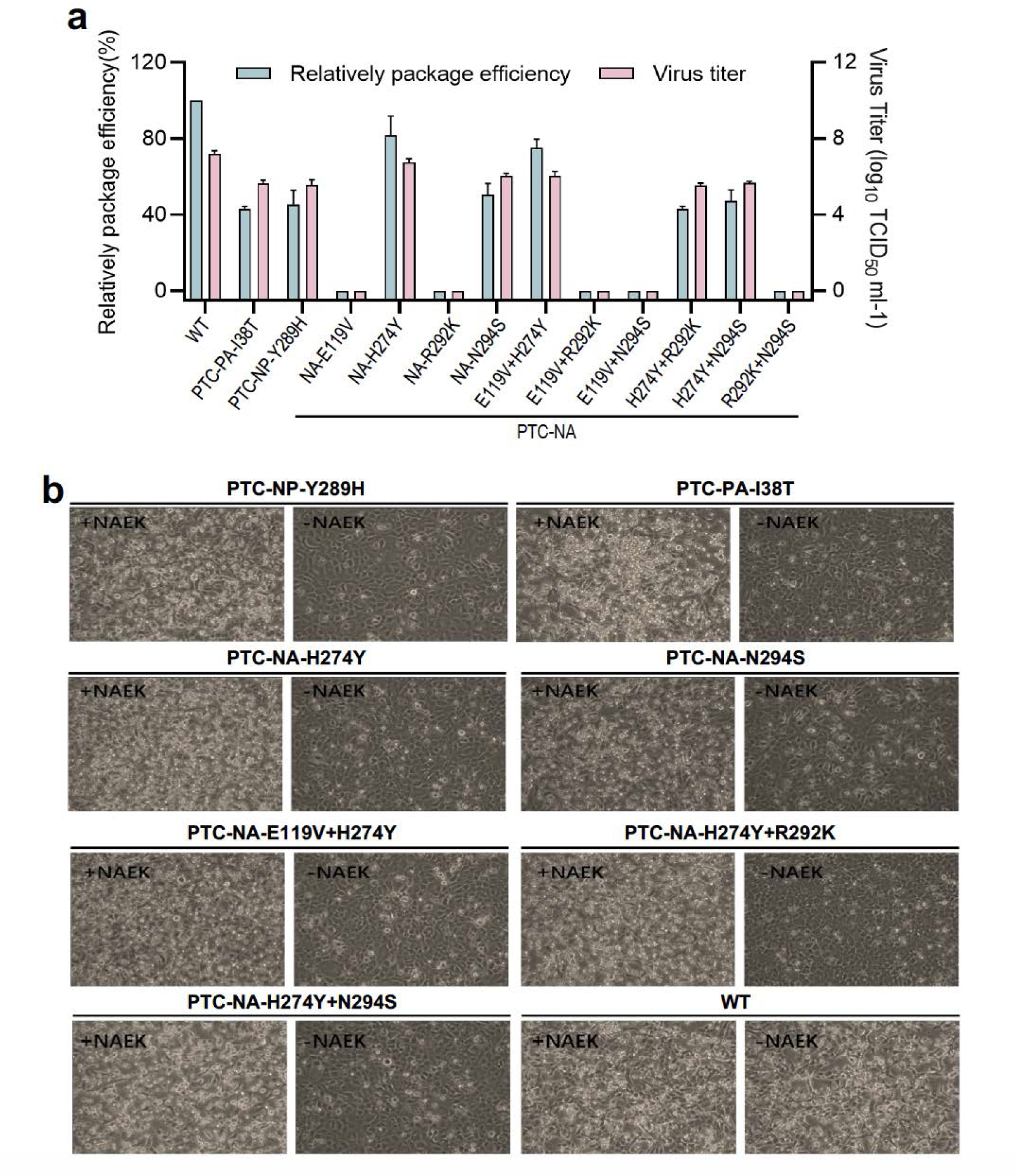
(**a**) Packaging efficiency and virus titers of different DR-UAA viruses. Data are presented as the mean ± SD (*n*=3). (**b**) The cytopathic effect (CPE) of different DR-UAA viruses and their dependence on NAEK (**Related to** **Fig. 1f**).

**Extended Data Figure 4.**
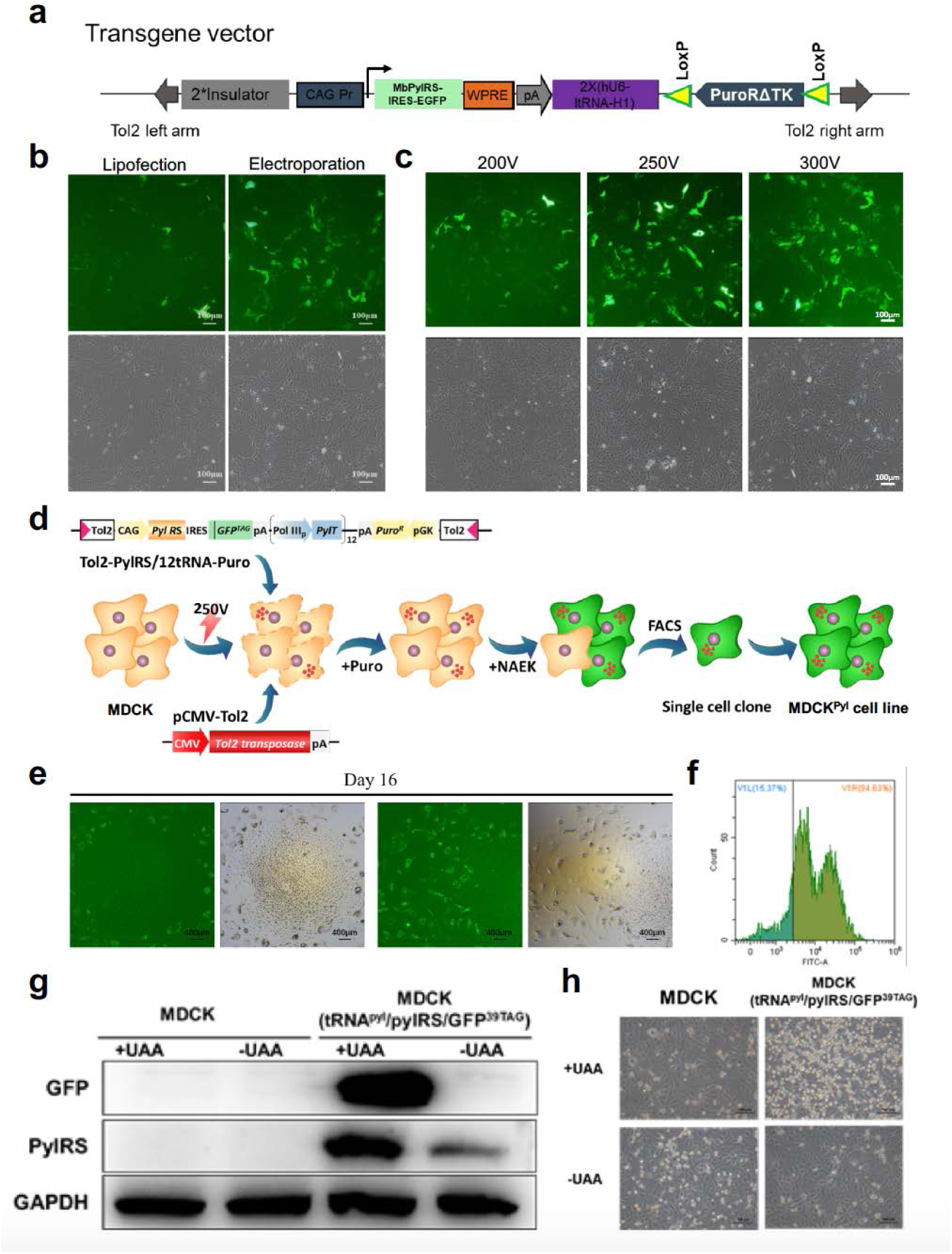
The construction of stable MDCK-tRNA^pyl^/PylRS/GFP^39TAG^ cell line. (**a**) The plasmid for the UAA incorporation system. (**b**) Comparison of transfection efficiency between lipofection and electroporation. (**c**) Comparison of transfection efficiency at different voltage. (**d**) Screening flow of MDCK-tRNA^pyl^/PylRS/GFP^39TAG^ stable cell line. (**e**) Positive clone for the MDCK-tRNA^pyl^/PylRS/GFP^39TAG^ stable cell line on day 16. (**f**) Fluorescent-activated cell sorting (FACS) for the MDCK-tRNA^pyl^/PylRS/GFP^39TAG^ stable cell line. (**g**) Western blot of green-fluorescent protein and PylRS expression. (**h**) The CPE of PTC virus and its dependence on UAA.

